# Reciprocal interactions between CA1 pyramidal and axo-axonic cells control sharp wave-ripple events

**DOI:** 10.1101/2024.07.02.601726

**Authors:** Earl T. Gilbert, Lianne M.F. Klaver, Kaiser C. Arndt, Jongwoon Kim, Xiaoting Jia, Sam McKenzie, Daniel Fine English

## Abstract

Diverse sources of inhibition serve to modulate circuits and control cell assembly spiking across various timescales. For example, in hippocampus area CA1 the competition between inhibition and excitation organizes spike timing of pyramidal cells (PYR) in network events, including sharp wave-ripples (SPW-R). Specific cellular-synaptic sources of inhibition in SPW-R remain unclear, as there are >20 types of GABAergic interneurons in CA1. Axo-axonic cells (AAC) are defined by their synaptic targeting of the axon initial segment of pyramidal cells, potently controlling spike output. The impact of AAC activity on SPW-R is controversial, due mainly to ambiguity of AAC identification. Here we monitored and manipulated opto-tagged AACs in behaving mice using silicon probe recordings. We found a large variability of AAC neurons, varying from enhanced to suppressed spiking during SPW-Rs, in contrast to the near-uniform excitation of other parvalbumin-expressing interneurons. AACs received convergent monosynaptic inputs from local pyramidal cell assemblies, which strongly influenced their participation in SPW-Rs. Optogenetic silencing of AACs increased power and duration of SPW-Rs, recruiting a greater number of PYR, suggesting AACs control SPW-R dynamics. We hypothesize that lateral inhibition by reciprocal PYR-AAC interactions thus supports the organization of cell assemblies in SPW-R.

## Introduction

Inhibitory actions in the cortex and hippocampus are diverse, with >20 types of inhibitory interneurons existing along a spectrum, with varied genetic profiles, axonal targeting, and state dependent activity^1–5^. Given the complexity that emerges from the nonlinear dynamics of inhibitory interactions among these types, numerous functions have been attributed to both their global and specific operations^6–8^. Studying unique interneuron subtypes in the hippocampus has proven fundamental in understanding mechanisms underlying circuit function, including their role in various network oscillations, learning, place field tuning and replay of experience^3,4,9–13^.

Axo-axonic cells (AACs) are anatomically distinct from other interneurons in two ways. First, only AACs target the axon initial segment of principal cells. Second, they do not target other interneuron types or each other, but instead, mutually interact with principal cells^14–16^. Despite the hypothesized importance of their distinct anatomical features, knowledge about AAC firing patterns in the behaving animal and their role in circuit control is limited^3,14,17,18^. Spiking activity of AACs has been mainly examined in connection with hippocampal sharp wave ripples (SPW-R), a network event critical in memory selection and consolidation, but their specific role has remained controversial^3,17–20^.

Leveraging the recent availability of the Unc-5b-Cre mouse-line^14^, which enables genetic access to AACs, we conducted *in vivo* silicon probe recordings to characterize the spiking activity of CA1 AACs in SPW-R. We report that participation of AACs in SPW-R falls along a continuum, driven by diverse interactions with CA1 pyramidal cells. Using optogenetic inhibition of AACs we found that AACs exhibit control over CA1 circuitry, most strongly during SPW-R. SPW-R that occurred during AAC-OFF epochs were of longer duration and higher power, though unchanged in frequency. Similarly, during AAC-OFF SPW-R, CA1 pyramidal cell recruitment was enhanced, suggesting that the magnitude of circuit activation during SPW-R, and potentially cell assembly selection, is controlled by AACs. These results demonstrate a critical role for AACs in CA1 SPW-R.

## Results

To characterize the activity of AACs, we recorded LFP and ensemble spiking in the hippocampal CA1 region of awake mice expressing excitatory or inhibitory opsins in AACs (**Fig 1A**). Selective opsin expression was achieved by injecting AAV2/5 encoding Cre-dependent opsins into the CA1 layer of adult Unc5B-CreER mice^14^, allowing us to opto-tag AACs. AACs were identified by activation or inactivation (ChR2 or Arch expressing AACs; **Fig 1B**). We identified cellular-synaptic properties of 43 AACs and examined how their reciprocal interactions with CA1 pyramidal cells organize circuit activity during SPW-R.

**Figure 1:**
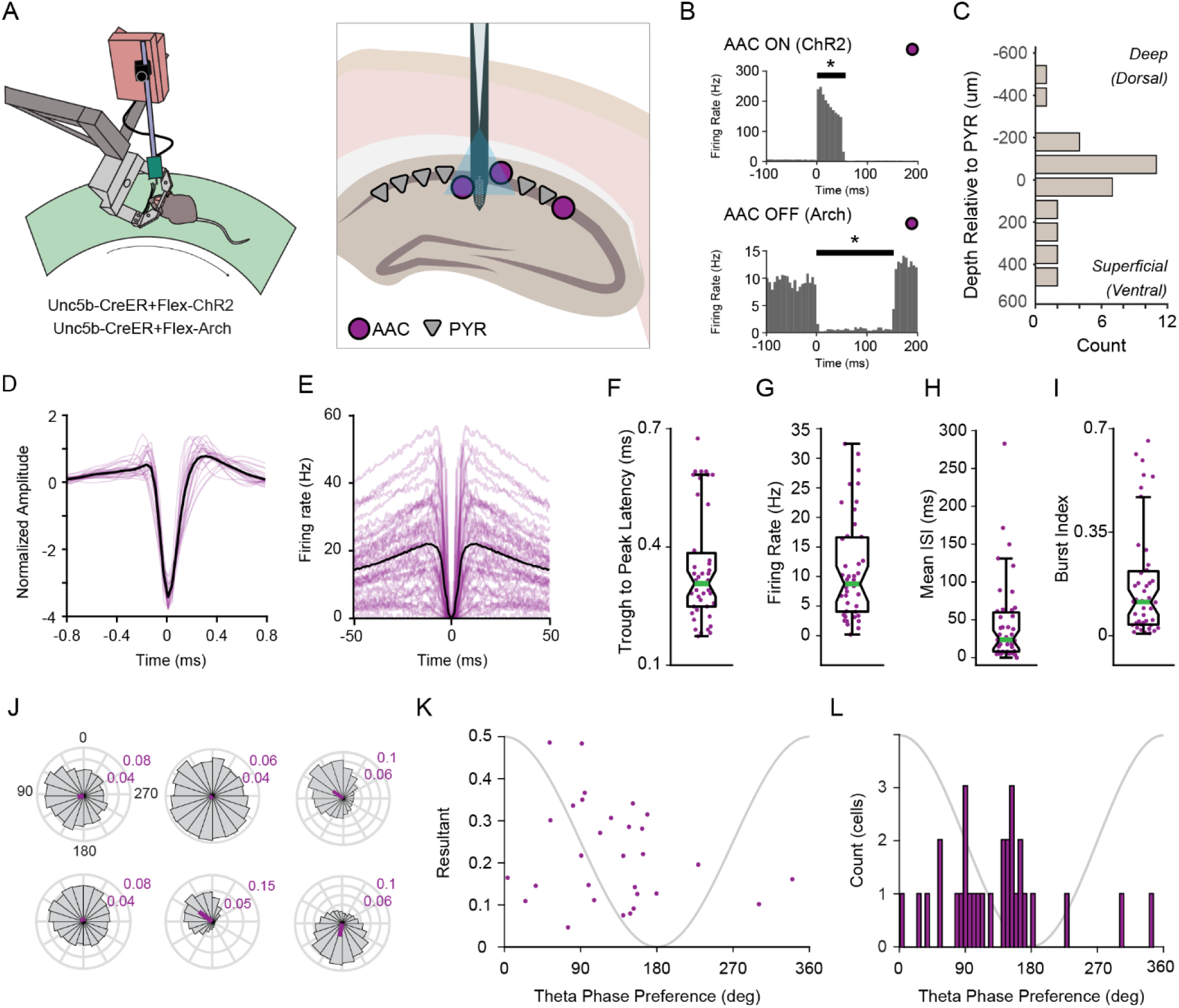
Firing characteristics of axo-axonic cells. A.) Cartoon of experimental set-up. Head-fixed mice run on a disc with an acutely implanted silicon probe in CA1 of the hippocampus. A sharpened optic fiber, glued to the probe, is used to stimulate axo-axonic cells. B.) Example peri-stimulus time histograms of two units’ responses to ChR2 (top) or Arch (bottom) activation. Putative interneurons with significant responses to the stimulus were identified as opto-tagged AACs. C.) Soma position of AACs relative to the pyramidal layer of CA1 (n=32 cells, µ=6.0µm, σ=215.6) D.) Average (black) and individual (purple) mean extracellular spike waveforms (n=43 cells) E.) Average (black) and individual (purple) spike autocorrelations (n=43 cells) F.) Trough to peak latency of extracellular waveforms. (n=43 cells, µ=0.35, σ=0.14) G.) Average firing rate (Hz) of AACs (n=43 cells, µ=11.14Hz, σ=8.62) H.) AAC inter-spike-interval (ms, n=43 cells, µ=45.45ms, σ=55.95) I.) Distribution of burst index (n=43 cells, µ=0.1747, σ=0.1906) J.) Example polar plots of spike entrainment to the theta oscillation of 6 AACs from one recording session. K.) Theta phase modulation versus phase preference of significantly theta modulated AACs (n=30 cells, Circular-Linear correlation, rho=0.7872, p=9.1922e-05) L.) Distribution of significantly theta modulated AACs phase preferences (n=30 cells)

### Characteristics of CA1 axo-axonic cells

We quantified physiological metrics of AACs (n=43 AACs, n=7 animals, 17 sessions). The somata of AACs were observed mostly in or close to the pyramidal layer (**Fig 1C**, µ=4.33µm relative to center of pyramidal layer, σ= 212.4) in agreement with previous studies^14,17,19^. Spike width was variable, (**Fig 1D**) with a trough to peak latency range of 0.18-0.68ms (**Fig 1F**, µ=0.35ms, σ=0.14), as were spike autocorrelations (**Fig 1E**). Mean firing rates were observed from 0.18-32.47hz (**Fig 1G**, µ=11.14hz, σ=8.62), with inter-spike intervals (ISI) ranging between 0.41ms to 285.71ms (**Fig 1H**, µ=45.45ms, σ=55.95). A small group of AACs showed transiently fast spiking (“bursting”), while the population had an observed range of 0.0036-0.6621 (burst index is fraction of spikes with less than 6-ms ISI^21^; **Fig 1I**, µ=0.1747, σ= 0.1906). This unexpected spectrum of physiology is interesting as it stands in contrast to the notion that AACs are monolithic, rather they exhibit significant intra-class variability as suggested in Varga et al., 2014^17^.

### Axo-axonic cells preferentially spike at the descending phase of the local theta oscillation

Prior reports have shown strong modulation of AAC spiking by phase of the ongoing theta rhythm^3,17^. Here we report in the largest to date collection of AACs (n=43). We found that many AACs were strongly theta modulated, similar to many CA1 interneurons. Theta phase preference of six AACs from a single recording session are shown in **Fig 1J**, demonstrating the observed variability is not due to inter-animal differences. The majority of AACs exhibit significant theta phase modulation (30/32, **Supplementary Table 1,** Rayleigh z test). Theta phase preference and phase modulation were correlated (**Fig 1K**, rho=0.7872, p=9.1922e-5, Circular-linear correlation coefficient), and most AACs spikes were distributed on the descending phase of theta(**Fig 1L**). The average theta phase preference for the population was 110.28°, while the most theta modulated AAC exhibited a phase preference of 91.04°. AACs as a population are highly theta modulated cells, with a phase preference for the descending phase of the oscillation^3,17,18^.

### Axo-axonic cells are activated in SPW-R

The participation of AACs in CA1 SPW-R is critical to understanding their role in the CA1 circuit, though there are mixed reports as to their level of participation^3,14,17–20^. Ripple-associated spiking activity of 43 AACs (**Fig 2A**, n=43 cells, n=595-16,058, µ=5,666.4 SPW-R per recording session), revealed diverse participation across the population. On average AACs have significantly increased firing rate during SPW-R (**Fig 2B**, Left: n=43 cells, baseline 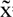=6.13hz, SPW-R 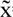=10.69hz, p=3.5132e-4, Wilcoxon signed-rank test). Interestingly, out of the 40 AACs that changed their firing rate (ZETA^22^), 27 cells increased and 13 cells decreased their activity (**Fig 2B**, Right), which stands in contrast to other types of interneurons with a more uniform response, such as PV+ cells (**Fig S1,** 43/46 cells showed increased activity). AACs demonstrated strong ripple phase modulation with most cells firing on the ascending phase of the ripple oscillation (100-250Hz, **Fig 2C, θ**_µ_=255.02 degrees), similar to previous reports for PV+ cells.

**Figure 2:**
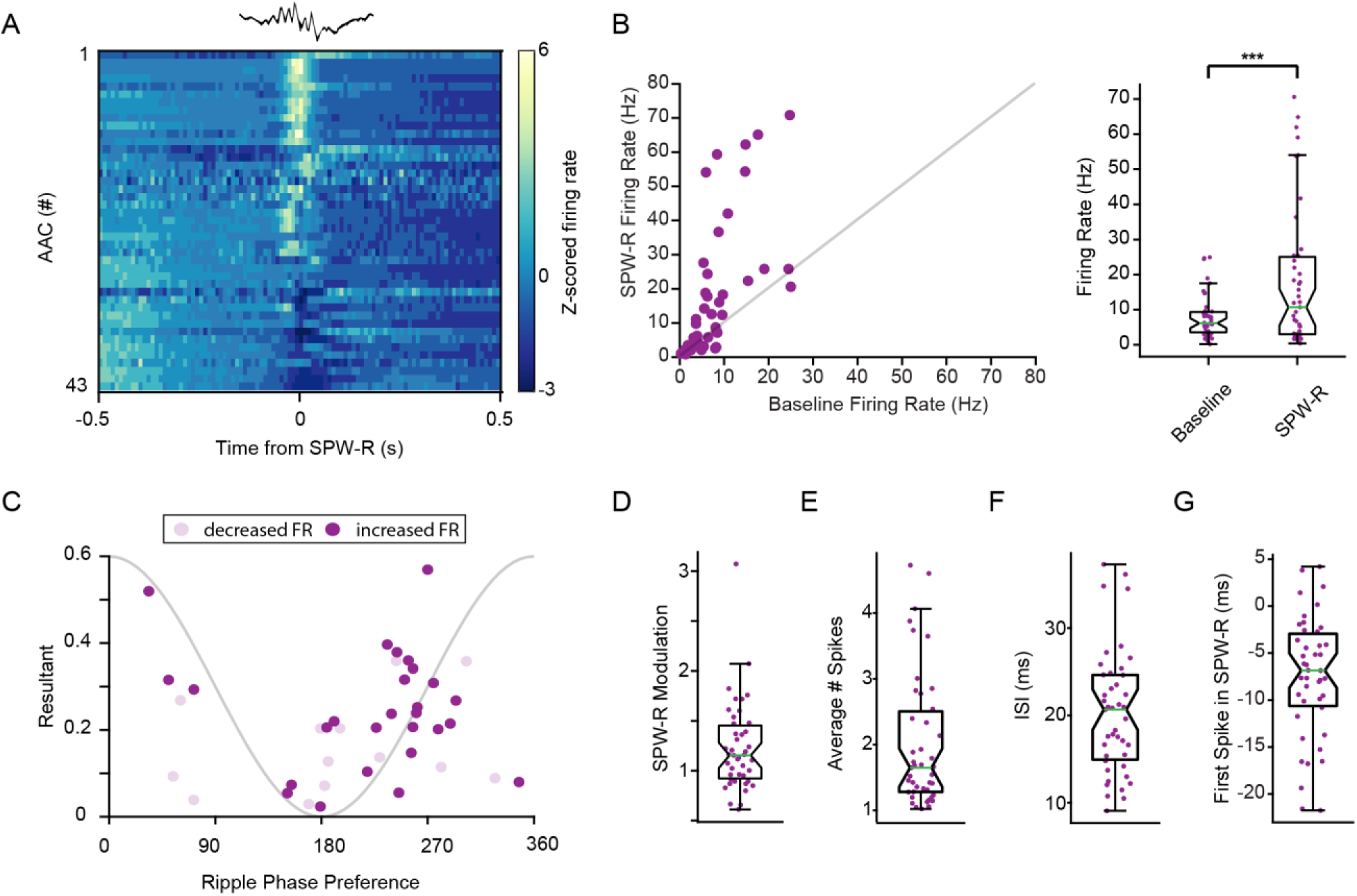
AACs firing is modulated by sharp wave-ripples. A.) Peri-event time histogram of AAC z-scored firing rate centered on SPW-R peak (n=43 cells, 1,495-16,058 SPW-R events, µ=5,666.4 events) B.) Left: Scatter plot of baseline firing rate versus firing rate during SPW-R of each AAC. (n=43). Right: Firing rate (Hz) in baseline and within SPW-R. (n=43 cells, baseline 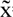=6.13hz, SPW-R 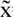=10.69hz, p=3.5132e-4, Wilcoxon signed rank test) C.) Ripple wave phase modulation versus phase preference of significantly SPW-R modulated AACs (n=40 cells) D.) SPW-R modulation of significantly modulated AACs (n=40 cells, µ=1.24, σ= 0.46) E.) Average number of spikes fired in SPW-R events (n=43 cells, µ=2.02, σ= 1.02) F.) Inter-spike-interval during SPW-R (n=43 cells, µ=20.3ms, σ= 7.0) G.) Average delay to first spike in SPW-R from the peak of SPW-R (n=43 cells, µ=-7.5ms, σ=6.4)

Single cell ripple modulation values (SPW-R firing rate gain) ranged from .61-3.01 (**Fig 2D**, µ=1.24, σ=0.46). The average number of spikes in SPW-R ranged from 1.02-4.73 (**Fig 2E**, µ=2.02, σ=1.02), while inter-spike intervals of AACs in SPW-R ranged from 9.1-37.2ms, approximating a range of 1-5 cycles of the oscillation (**Fig 2F**, µ=20.3ms, σ=7.0). Additionally, because SPW-R information content systematically varies from the beginning to end of each event, we calculated the average time of the first spike from the peak of the SPW-R, which ranged from −21.7-42.0ms (**Fig 2G**, µ=-7.5ms, σ=0.0064), suggesting AACs influence activity throughout the SPW-R event. Together these findings demonstrate that AAC activity in SPW-Rs is variable, however most AACs increase their activity in SPW-R events.

### CA1 pyramidal neurons control the participation of axo-axonic cells in SPW-R

To determine if local excitation is a mechanism by which AAC activity is modulated in SPW-R, we quantified excitatory synaptic inputs from local CA1 PYR. Such monosynaptic connections were identified using a validated monosynaptic inference method^23,24^ (based on cross-correlations of putatively-classified pyramidal cells and AACs). For AACs with at least one presynaptic PYR, we calculated the average SPW-R modulation of the presynaptic inputs. Example SPW-R cross-correlations (**Fig 3A**) illustrate both strongly and weakly modulated AACs and their partners (note that an AAC with a large peak in the cross-correlation at the time of SPW-R has partners that also show a large peak, **Fig 3A, top**). AAC ripple modulation correlated with that of its inputs, such that AACs that received synaptic drive from ripple modulated PYR were themselves strongly ripple modulated. (**Fig 3B**, r=0.6923, p=1.3325e-4, Spearman’s Rank Correlation; correlation with randomly selected non-presynaptic partner control, r=0.1915 p=0.3574, Spearman’s Rank Correlation; see Fig S2 for correlations with other physiological metrics.). We found further evidence that the CA1 PYR modulate AAC activity in SPW-R when examining other metrics. We quantified the correlation between spiking activity and power in the ripple frequency band as well as the magnitude of phase modulation. There was a significant relationship between spike rate power correlation of presynaptic PYR and AACs (**Fig 3C**, r=0.4619, p=0.0185, Spearman’s Rank Correlation), as well as ripple phase modulation between paired cells (**Fig 3D** r=0.5015 p=0.0099, Spearman’s Rank Correlation). Given the relationship between presynaptic and AAC activity, we investigated the relationship between strength of presynaptic connections and AAC participation in SPW-R. Both AAC SPW-R modulation and time to first spike were related to spike transmission probability. Stronger spike transmission probability was associated with higher ripple modulation (**Fig 3E**, r=0.5508, p=0.0041, Spearman’s Rank Correlation), and a shorter delay between first spike and SPW-R peak (**Fig 3F**, r=-0.7039, p= 9.2163e-5, Spearman’s Rank Correlation). These results suggest that excitation from CA1 PYR controls AAC activity during SPW-R.

**Figure 3:**
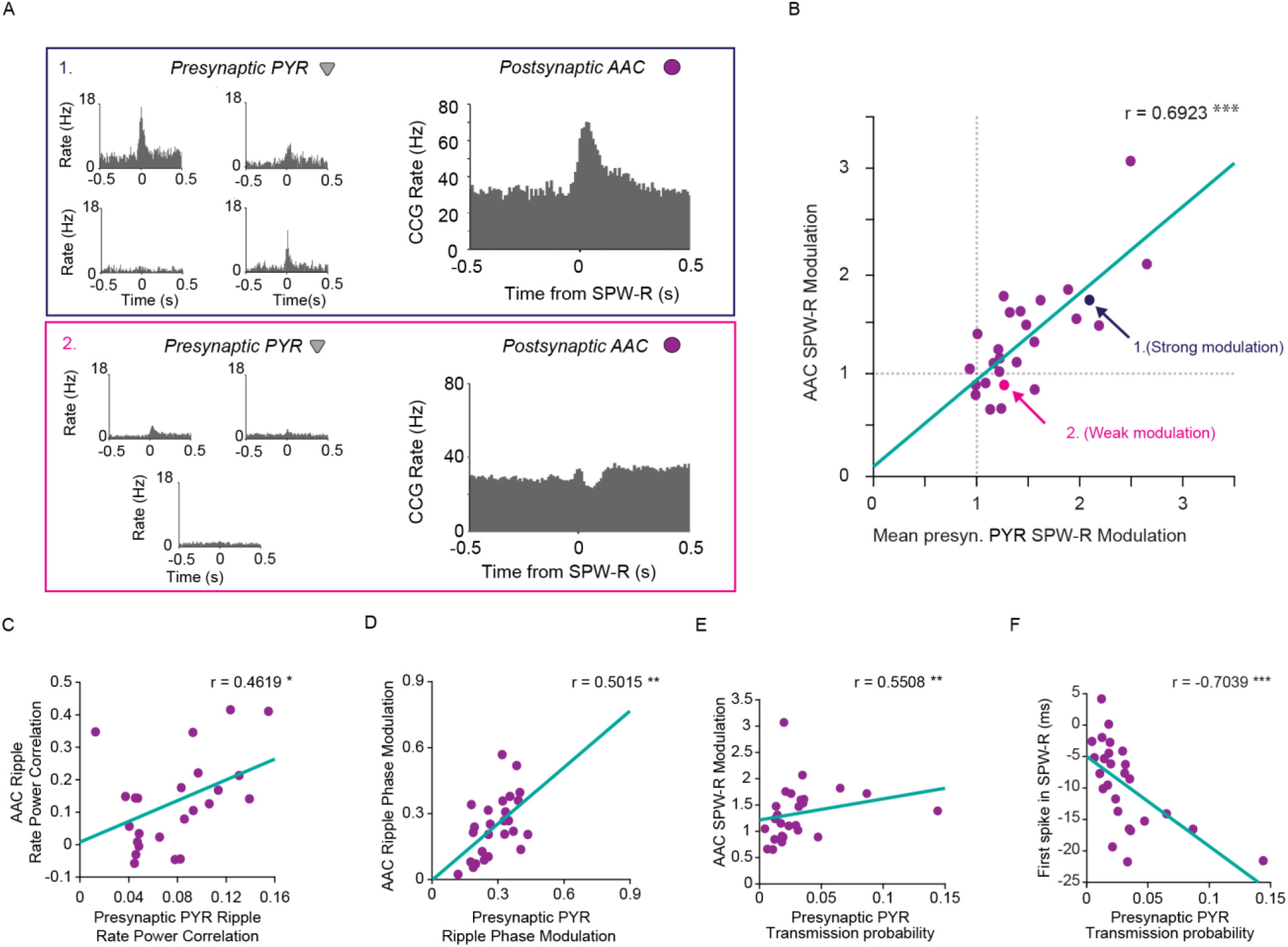
Local presynaptic pyramidal cells contribute to spiking of AACs in SPW-R. A.) Top: SPW-R cross-correlograms between 4 pyramidal cells presynaptically (prePYR) connected to a strongly ripple-activated AAC (right). Bottom: Example SPW-R cross-correlograms between 3 pyramidal cells presynaptically connected to a of a weakly ripple modulated AAC. B.) Correlation between average SPW-R modulation of presynaptic PYR and their monosynaptically connected AACs (n=25 AACs, n=1-8 prePYR, r=0.6923, p=1.3325e-4, Spearman’s Rank Correlation). Arrows point to the example sets in A. C.) Correlation between average rate power correlations of spike to ripple frequency (100-250Hz) power of presynaptic partners and AACs (n=25 AACs, n=1-8 prePYR, r=0.4619, p=0.0185, Spearman’s Rank Correlation) D.) Correlation between average ripple (100-250Hz) phase modulation of presynaptic partners and AACs (n=25 AACs, n=1-8 prePYR, r=0.5015 p=0.0099, Spearman’s Rank Correlation) E.) Correlation between average PYR-AAC spike transmission probability and SPW-R modulation of AACs (n=25 AACs, n=1-8 prePYR, r=0.5508, p=0.0041, Spearman’s Rank Correlation) F.) Correlation between average PYR-AAC spike transmission probability and AAC average delay to first spike in SPW-R from peak (n=25 AACs, n=1-8 prePYR, r=-0.7039, p= 9.2163e-5, Spearman’s Rank Correlation)

### AACs control network-wide spiking activity and SPW-R dynamics

We found that AACs robustly participate in SPW-R, though whether they influence the generation of SPW-R in the network remains unknown. To address this question, we optogenetically inhibited local AACs expressing archaerhodopsin (Arch). Silencing AACs increased power in the ripple band (100-250hz, **Fig 4A,B**, Bottom: n=5 sessions, 4 mice. n=843-1,936 300ms stims. p=0.0020, Paired T-test), as well as power in the 300-500Hz band, which reflects population spiking activity (300-500hz, **Fig 4A,B**, Top: n=5 sessions, 4 mice. n=843-1,936 300ms stims. P=3.0224e-04, Paired T-test). We then detected SPW-R and examined properties of such events occurring either in or out of AAC inhibition epochs. Subtraction of baseline ripple PSDs from AAC-OFF ripple PSDs reveal increased power of the ripple frequency at the time of these events. (**Fig 4C**, n=5 sessions, 4 mice. n=1,495-12,364 baseline ripples, n= 64-335 stim ripples), as did event durations (**Fig 4E**, n=5 sessions, 4 mice. n=37,032 baseline ripples, n= 1,024 stim ripples. p=0.0072, Wilcoxon rank sum-test), and peak amplitude (**Fig 4F**, n=5 sessions, 4 mice. n=37,032 baseline ripples, n=1,024 stim ripples. p=2.0593e-06, Wilcoxon rank sum test), while ripple frequency was not affected (**Fig 4D**, n=5 sessions, 4 mice. n=37,032 baseline ripples, n=1,024 stim ripples. p=0.8675, Wilcoxon rank-sum test). This suggests that AACs control content and magnitude of SPW-R while other cell types may pace the rhythm (likely basket cells^25^). These data reveal an important role for AACs in controlling SPW-R dynamics at the network level.

**Figure 4:**
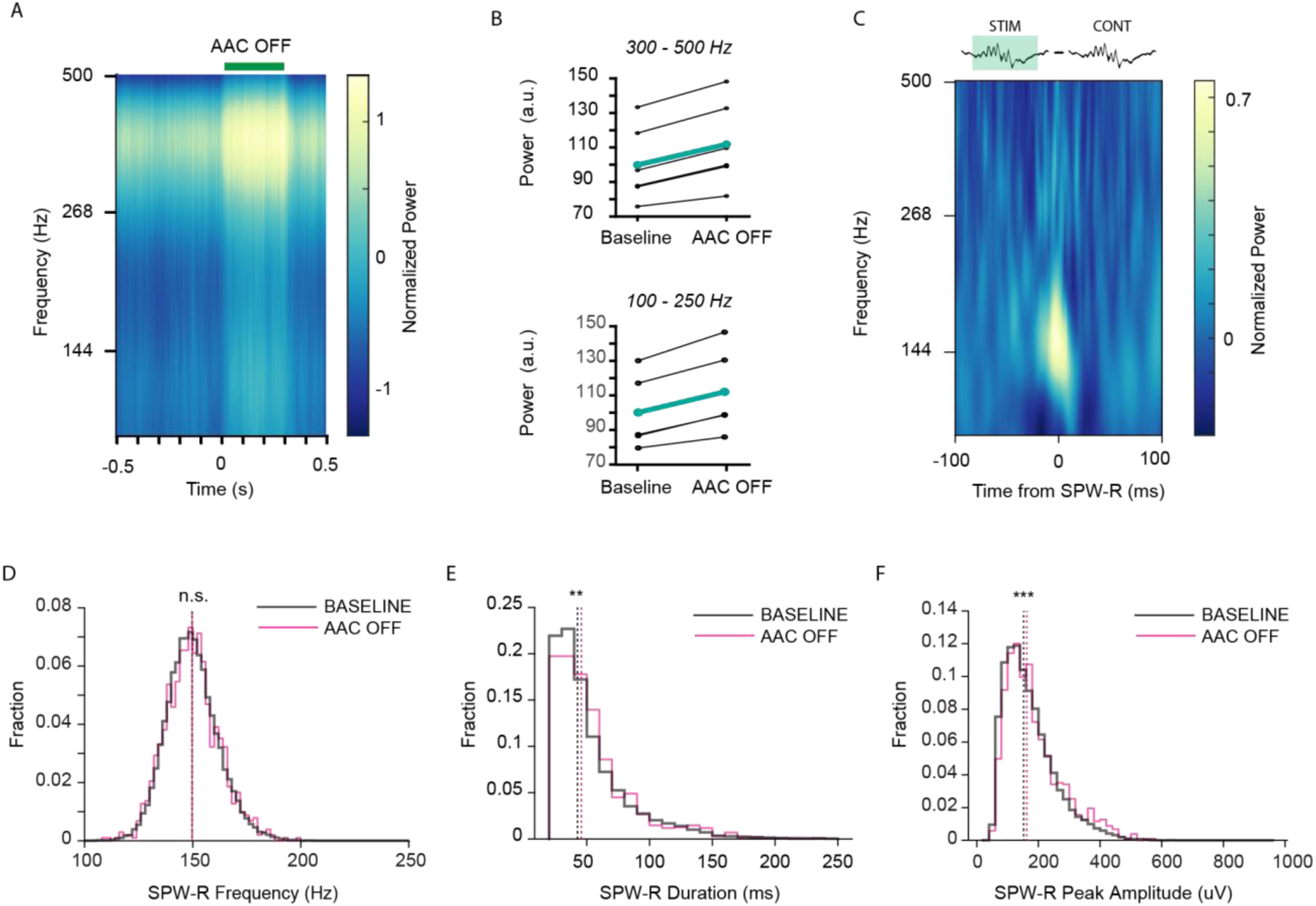
Axo-axonic cells affect SPW-R dynamics. A.) Mean peri-stimulus power spectral density (PSD) of AAC-Arch activation (n=5 sessions, 4 mice. n=843-1,936 300ms stims) B.) Average power of PSD in A comparing baseline (−300ms:0ms) to stimulation (0ms:300ms) in the spike band (top, 300-500Hz n=5 sessions, 4 mice. n=843-1,936 300ms stims. p=3.0224e-04, Paired T-test) and the SPW-R band (bottom, 100-250Hz. n= 5 sessions, 4 mice. 843-1,936 300ms stims. p=0.0020, Paired T-test) C.) Average SPW-R event PSD of baseline events subtracted from AAC-OFF events (n=5 sessions, 4 mice. n=1,495-12,364 baseline ripples, n=64-335 AAC-OFF ripples.) D.) Distribution of frequency of AAC-OFF SPW-R events compared to baseline control (n=5 sessions, 4 mice. n=37,032 baseline ripples, n= 1,024 AAC-OFF ripples. p=0.8675, Wilcoxon rank sum test.) E.) Distribution of duration of AAC-OFF SPW-R events compared to baseline control (n=5 sessions, 4 mice. n=37,032 baseline ripples, n= 1,024 AAC-OFF ripples. p=0.0072, Wilcoxon rank sum test.) F.) Distribution of peak amplitude of AAC-OFF SPW-R events compared to baseline control (n=5 sessions, 4 mice. n=37,032 baseline ripples, n= 1,024 AAC-OFF ripples. p=2.0593e-06, Wilcoxon rank sum test.)

### AACs control pyramidal cell participation in SPW-R

Pyramidal cells are reliably depolarized during SPW-R, and competition between excitatory and inhibitory inputs compete to determine spiking at the single cell level^12^. Given our findings that most AACs are active in SPW-R, and inhibiting them alters SPW-R dynamics, we next investigated the effects of silencing AACs on PYR spiking. We first looked at the effects of manipulating AAC activity in all states. AAC activation inhibited PYR firing, while AAC silencing resulted in PYR disinhibition, revealed by changes in their firing rates. (**Fig 5A**, Left, ChR2: n=11 sessions, 4 mice. n=1,063-4,459 100ms stims, n=180 PYR, Right, Arch: n=5 sessions, 4 mice. n=843-1,936 300ms stims, n=207 PYR). These results demonstrate that AACs tonically inhibit PYR. Next, we sought to investigate how AAC inhibition shapes PYR activity selectively during SPW-R by comparing baseline ripples to ripples that occurred during periods where AACs were silenced. We quantified the fraction of PYR that participated in AAC-OFF SPW-R compared to baseline. In AAC-OFF SPW-R, we observed a significant increase in PYR participation (**Fig 5B**, Left: AAC: n=5 sessions, 4 mice. n=31-55, total: 194 PYR. n=1,495-12,364, total: 37,032 Baseline SPW-R. n=51-287, total: 868 AAC-OFF SPW-R. p=5.2716e-16, Wilcoxon rank-sum test). Superficial and deep CA1 PYR receive different patterns of inhibition^26,27^, thus we quantified this effect separately for each subgroup of pyramidal population. In AAC-OFF SPW-R, the fraction of both superficial and deep PYR participation increased (**Fig S3**, Right: Top, AAC, n=1,495-12,364, total: 37,032 Baseline SPW-R. n=51-287, total: 868 AAC-OFF SPW-R. n=11-32, total: 125 Deep cells, n=5-33, total: 69 Superficial cells. Deep: p=2.4645e-13, Wilcoxon rank-sum test. Superficial: p=6.5141e-06, Wilcoxon rank-sum test.). Given that cell assemblies constructed from unique sets of PYR enable the encoding of separate experiences by SPW-R^28–31^ we examined SPW-R associated PYR activity at an individual cell level. On average, PYR exhibited a higher count of spikes in AAC-OFF SPW-R (**Fig 5C**, n=5 sessions, 4 mice. n=194 PYR, n=1,495-12,364 Baseline SPW-R. n=51-287 AAC-OFF SPW-R. p=6.1155e-08, Wilcoxon signed-rank test). Similarly, pyramidal cell SPW-R firing rate gain was significantly higher when AACs were silenced (**Fig 5D**, n=5 sessions, 4 mice. n=194 PYR, n=1,495-12,364 Baseline SPW-R. n=51-287 AAC-OFF SPW-R. p=2.4569e-08, Wilcoxon signed-rank test). Increased firing rate was naturally reflected in decreased inter-spike interval in AAC-OFF SPW-R. (**Fig 5E**, n=5 sessions, 4 mice. n=194 PYR, n=1,495-12,364 Baseline SPW-R. n=51-287 AAC-OFF SPW-R. p=6.9834e-05, Wilcoxon signed-rank test.) Further, AACs participated in a higher fraction of total AAC-OFF SPW-R than baseline SPW-R. (**Fig 5F**, n=5 sessions, 4 mice. n=194 PYR, n=1,495-12,364 Baseline SPW-R. n=51-287 AAC-OFF SPW-R. p=1.8104e-11, Wilcoxon signed-rank test). These results demonstrate AACs control the participation of pyramidal cells in SPW-R events, likely contributing to functions such as replay.

**Figure 5:**
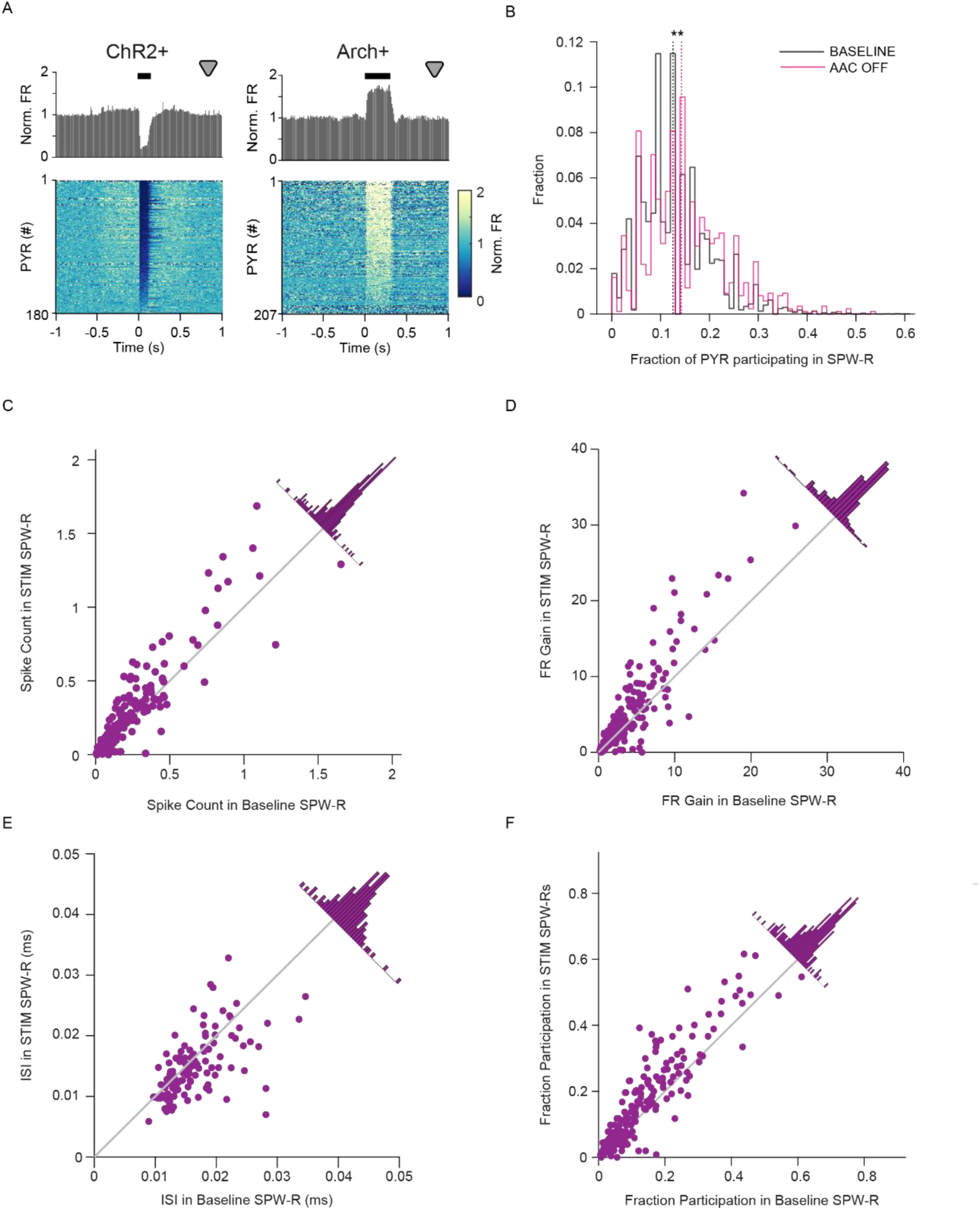
AACs control pyramidal cell participation during SPW-R. A.) Peri-stimulus time histograms of PYR firing rate in response to stimulation of interneurons in a subset of sessions with matched stim durations. (**Left**, ChR2: n=11 sessions, 4 mice. n=1,063-4,459 100ms stims, n=180 PYR, **Right**, Arch: n=5 sessions, 4 mice. n=843-1,936 300ms stims, n=207 PYR) B.) Distribution of fraction of PYR participating in baseline and AAC-OFF SPW-R. (n=5 sessions, 4 mice. n=31-55, total: 194 PYR. n=1,495-12,364, total: 37,032 Baseline SPW-R. n=51-287, total: 868 AAC-OFF SPW-R. p=5.2716e-16, Wilcoxon rank-sum test.) C.) Scatter plot of individual PYR mean spike count in baseline SPW-R versus count in AAC-OFF SPW-R. Gray: Unity Line. Distribution of distances from unity line on diagonal. (n=5 sessions, 4 mice. n=194 PYR, n=1,495-12,364 Baseline SPW-R. n=51-287 AAC-OFF SPW-R. p=6.1155e-08, Wilcoxon signed-rank test) D.) Scatter plot of individual PYR firing rate gain in baseline SPW-R versus firing rate gain in AAC-OFF SPW-R. Gray: Unity Line. Distribution of distances from unity line on diagonal. n=5 sessions, 4 mice. n=194 PYR, n=1,495-12,364 Baseline SPW-R. n=51-287 AAC-OFF SPW-R. p=2.4569e-08, Wilcoxon signed-rank test) E.) Scatter plot of individual PYR inter-spike interval in baseline SPW-R versus inter-spike interval in AAC-OFF SPW-R. Gray: Unity Line. Distribution of distances from unity line on diagonal. (n=5 sessions, 4 mice. n=122 PYR, n=1,495-12,364 Baseline SPW-R. n=51-287 AAC-OFF SPW-R. p=6.9834e-05, Wilcoxon signed-rank test.) F.) Scatter plot of fraction of baseline SPW-R in which individual PYR participated versus participation in AAC-OFF SPW-R. Gray: Unity Line. Distribution of distances from unity line on diagonal. (n=5 sessions, 4 mice. n=194 PYR, n=1,495-12,364 Baseline SPW-R. n=51-287 AAC-OFF SPW-R. p=1.8104e-11, Wilcoxon signed-rank test.)

## Discussion

Glutamatergic excitation and GABAergic inhibition compete to control PYR participation in SPW-R^25^. In each SPW-R, only a minority of PYR spike even though Vm is depolarized above spike threshold for 10’s of milliseconds, due to dynamically balanced GABAergic inhibition. Inhibition thus plays a critical role in enabling competition-based selection and expression of PYR assemblies. Identifying the source(s) of this inhibition is critical to understanding the CA1 circuit, and likely arises from axo-somatic targeting cells such as basket cells and AACs. Here we identify novel mechanisms by which AACs participate in and control CA1 SPW-R. Notably, these findings strongly suggest that reciprocal interactions between excitatory PYR and inhibitory AACs organize cell assembly construction and overall network activity during SPW-R. This places AACs in a critical position in determining the content of CA1 information delivered to neocortex.

### Recruitment of AACs to sharp wave-ripples is explained by monosynaptic inputs from CA1 PYR

CA1 interneuron spiking is controlled both by afferent inputs (e.g., CA3) and excitation and inhibition from local PYR and INT, respectively^32–34^. To determine the activity of AACs and PYR during SPW-R at single-spike resolution, we used silicon probes to obtain CA1 ensemble recordings and opto-tagged AACs. As a population, AAC activity is significantly increased during SPW-R, with a minority of cells exhibiting decreased activity, emphasizing the importance of considering intra-class variability in studying cell types^17^ (**Fig 2A,B)**. In agreement with these findings, up- and down-modulated AACs have been previously described in studies using both juxta-cellular electrophysiology^26^ and calcium imaging^14,19^. To better understand the driver of AAC spiking in SPW-R, we examined their physiology and excitatory inputs from CA1 PYR. Individual AAC activity in SPW-R is strongly correlated with the activity of presynaptic partners and the strength of these connections (**Fig 3)**, suggesting that the local circuit can control AAC activity independently of afferent input. An important future question is if the propensity for AACs to participate in SPW-R is consistent across experiences and/or throughout the lifespan of the animal^35^. Such plasticity could be supported both by changes in excitatory inputs we identified as well as inhibition from local interneurons^36^. Longitudinal recording of single AACs would be required to answer these questions.

### AACs control network activation and PYR spiking in sharp wave-ripples

Silencing AACs led to increased power in both the spiking band and SPW-R band (**Fig 4A,B**), SPW-R events that occurred during AAC silencing were of longer duration, and had higher power and peak amplitude (**Fig 4C,F,G**). Interestingly AAC silencing did not alter SPW-R frequency. These data support a model in which AACs robustly control PYR recruitment, while other interneurons such as PV+ cells may be more important for synchronizing PYR populations at ripple frequency^25^.

PYR activity in SPW-R was affected significantly by silencing AACs. Inhibition of AACs increased the number of participating PYR (**Fig 5B)** and produced increased average spike count, firing rate gain and decreased inter-spike intervals (**Fig 5C,D,E)**. Additionally, PYR participated in a greater fraction of AAC-OFF epochs than during control SPW-R, suggesting AACs function to select which PYR are allowed to participate in each event. Our findings that during SPW-R, silencing AACs disinhibits PYR, and that PYR drive AAC spiking, suggests lateral inhibition is instantiated by reciprocal interactions between pyramidal cells and AACs. These results support the general hypothesis that individual interneurons serve unique functions in CA1 circuit computations, and that this mechanism supports the construction of and competition between cell assemblies. Our experiments demonstrate this specifically for the understudied AAC interneuron family.

## Methods

All protocols and experiments using animals were approved by the Virginia Tech (Blacksburg, VA, USA) Institutional Animal Care and Use Committee (IACUC). The Unc5b-CreER mice were obtained from Z. Josh Huang under a fully executed MTA, and either maintained on a homozygous C57BL/6J background or were bred with FVBN/J wild-type mice to generate heterozygous mice. Male and female adult mice (>12 weeks) were used in all experiments.

### AAV Injection

Mice are induced and maintained at a surgical plane of anesthesia with isoflurane while mounted in a stereotaxic frame. Hair is removed and scalp is disinfected. Bupivicaine is injected once under the scalp before revealing skull. A < 0.2 mm burr hole is made above area CA1 of the hippocampus in both hemispheres. (mm from bregma: −1.8, lateral: +/-1.5). Glass pipette containing AAV5-EF1a-DIO-hChR2 (H134R)-EYFP or AAV5-EF1a-DIO-eArch3.0-EYFP (UNC Gene Therapy Center – Vector Core) is lowered into CA1 using stereotaxic coordinates. 100nL of AAV (titer: 3.8-5.2×10^12^ GC/mL) is injected into tissue (1nLs^−1^) using a nanoliter injector (WPI: MICRO2T & 504127). Following infusion, the glass pipette is left to rest for 5 minutes before slowly removing from the brain. The scalp is then closed with Vetbond (3M).

### Tamoxifen injections

Mice were injected intraperitoneally with tamoxifen (100 mg/kg) dissolved in corn oil at 20 mg/ml on days 5,7, and 9 following AAV injection.

### Head-bar Implantation

Mice are induced and maintained at a surgical plane of anesthesia with isoflurane while mounted in a stereotaxic mount. Bupivicaine is injected under the scalp before it is removed. The skull is cleaned with 3% hydrogen peroxide, followed by an application of dental adhesive Optibond (Kerr Dental). A small burr hole is made above cerebellum, and a stainless-steel ground wire is inserted between the skull and the brain, parallel to the brain surface, then the wire is affixed to the skull with sterile dental acrylic. A headplate is positioned parallel to the skull above lambda, and is permanently fixed in place.

### Craniotomy

One day before recording, mice are induced and maintained at a surgical plane of anesthesia with isoflurane while mounted in the stereotaxic mount. A <1.0 mm burr hole is made above the hippocampus and the dura is removed. Kwik-cast (World Precision Instruments) is applied to the burr hole, with dental acrylic applied on top, to protect the craniotomy until recording.

### *In Vivo* recording

Unc5b-CreER mice were obtained from the lab of Z. Josh Huang under full MTA. All animals used in experiments were homozygous Unc5b-CreER or hybrid Unc5b-CreER x FVB/NJ (JAX #001800) mice. Additionally, PV-Cre mice were acquired from the Jackson Laboratory (Jax #017320). During the time of experiments, mice were between the ages of 16–30 weeks old. Mice of both sexes were used in all experiments. Mice, trained and habituated to head-fixed navigation, are placed in the head fixation apparatus, and a silicone probe (Poly3, Neuronexus, or P1, Cambridge Neurotech, with optic fiber attached, or 4×8 uLED, Neurolight technologies), is lowered through the craniotomy into the brain until SPW-R are observed and left in place for 30-45 minutes until tissue is relaxed. The neural signals were amplified, digitized and recorded with RHD2000 system (Intan Technologies LLC).

### Optical Stimulation

Optical pulses were delivered through a sharpened optic fiber connected to a Thorlabs laser diode driver and laser diode or optical pulses are delivered through LEDs in a 4×8 uLED probe (Neurolight Technologies).

### Data collection

Unc5b-CreER ChR2, Arch, and PV+ data was collected for this study. A selection of PV+ data was used with permission from the Buzsaki Data Repository^24,37^.

### Data analysis

Data analysis was conducted in Matlab (Mathworks) and custom scripts were used to analyze the extracellular and spike data. Spike sorting was performed using Kilosort1^38^ with custom semi-automated reclustering^39^ followed by manual curation using Phy. Cell Explorer^40^ was used to characterize neuron properties including cell position, autocorrelation, wave form characteristics, firing rate characteristics, putative monosynaptic connections (followed by manual refinement) and SPW-R modulation. The Buzcode repository was used to calculate cross correlations and for SPW-R detection. The HippoCookBook repository and Circular Statistics Toolbox (Mathworks) were used to quantify oscillation phase preference and phase modulation. Rate power correlation was calculated by correlating spike trains with power in the SPW-R band (100-250Hz). Opto-tagged AACs were identified using CellExplorer to classify putative cell type, and Zenith of Event-based Time-locked Anomalies (ZETA^22^) to determine significance of the response to stimulation. Further, opto-tagged AACs were manually refined through observation of peri-stimulation time histogram to optical stimulation. Significance of SPW-R modulation was determined using CellExplorer and ZETA. For detection of SPW-R characteristics in and out of stimulation (Fig 4), SPW-R were classified as having the peak of the event within stimulation times. For detection of spikes from cells in SPW-R events (Fig 5), only SPW-R that had the entire duration within a stimulation pulse were considered INT-OFF SPW-R.

## Acknowledgements

The research in this report was supported by grants to from the National Institute of Neurological Disorders and Stroke (NINDS) R01NS131858 (DFE and SM), The Whitehall Foundation (DFE) and The Simons Foundation (DFE). We would like to thank Gyorgy Buzsaki and Ivan Soltesz for comments on previous versions of this manuscript, Zachary Saccomano for assistance with analytical approaches, and Z. Josh Huang for providing the Unc5b-CreER mouse line.

## Author Contributions

E.G., L.M.F.K., S.M., and D.F.E. designed the study. E.G., S.M., L.M.F.K and D.F.E. conducted experiments. E.G., L.M.F.K., K.A., J.K., S.M., and D.F.E. conducted data analysis. All authors contributed to the writing of the manuscript.

## Supplemental Information

**Table 1: Quantification of AAC characteristics**

**Figure S1:**
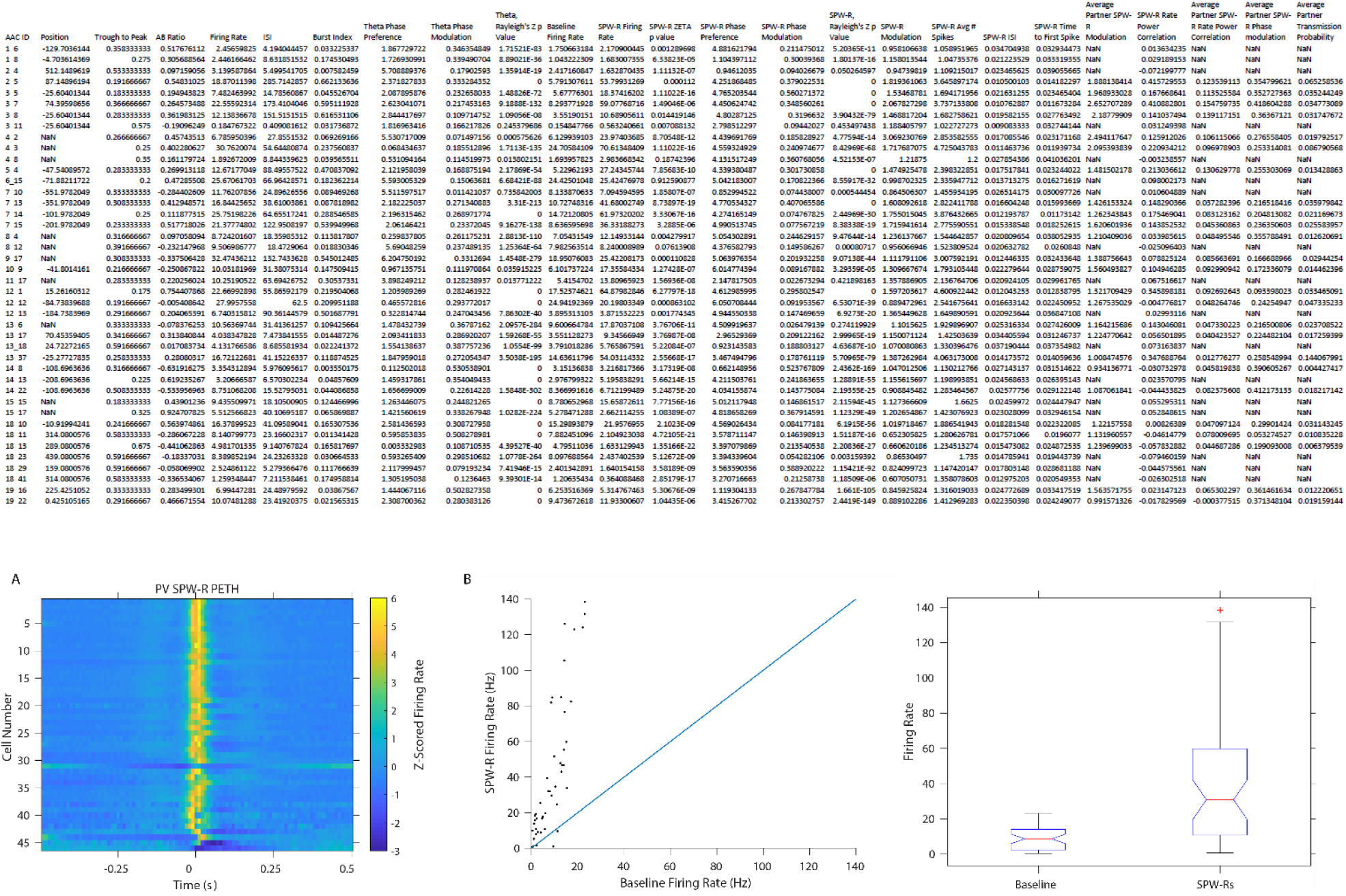
PV+ cells increase their firing rate during SPW-R. A.) Peri-event time histogram of PV z-scored firing rate centered on SPW-R peak (n=46 cells, n=6 animals, 11 sessions. 2,031-10,960 SPW-R events, µ= 6,725.6 events) B.) Left: Firing rate (Hz) in baseline and within SPW-R. (n=46 cells, n=6 animals, 11 sessions. baseline 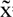= 8.63Hz, SPW-R 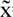= 30.82Hz, p=3.0939e-4, Wilcoxon signed rank test) Right: Distribution of delta firing rate in and out of SPW-R (n=46 cells).

**Figure S2:**
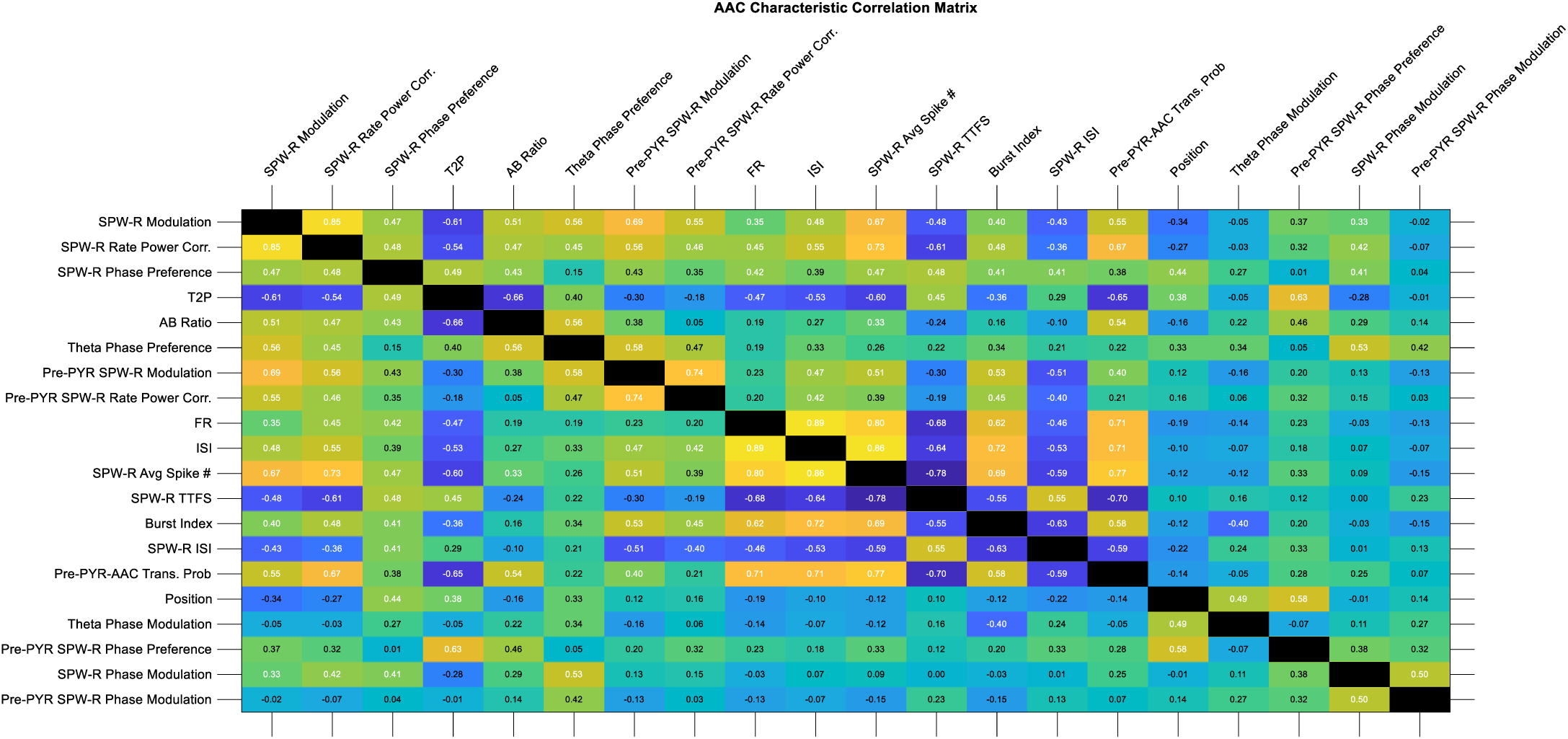
Correlation Matrix of AAC characteristics. Clustered correlation matrix of AAC characteristics including SPW-R metrics of presynaptic PYRs. Significant correlations are labeled in white text (p<.05, Spearman’s Rho, Circular-Linear Correlation Coefficient, and Circular-Circular Correlation Coefficient, respectively).

**Figure S3:**
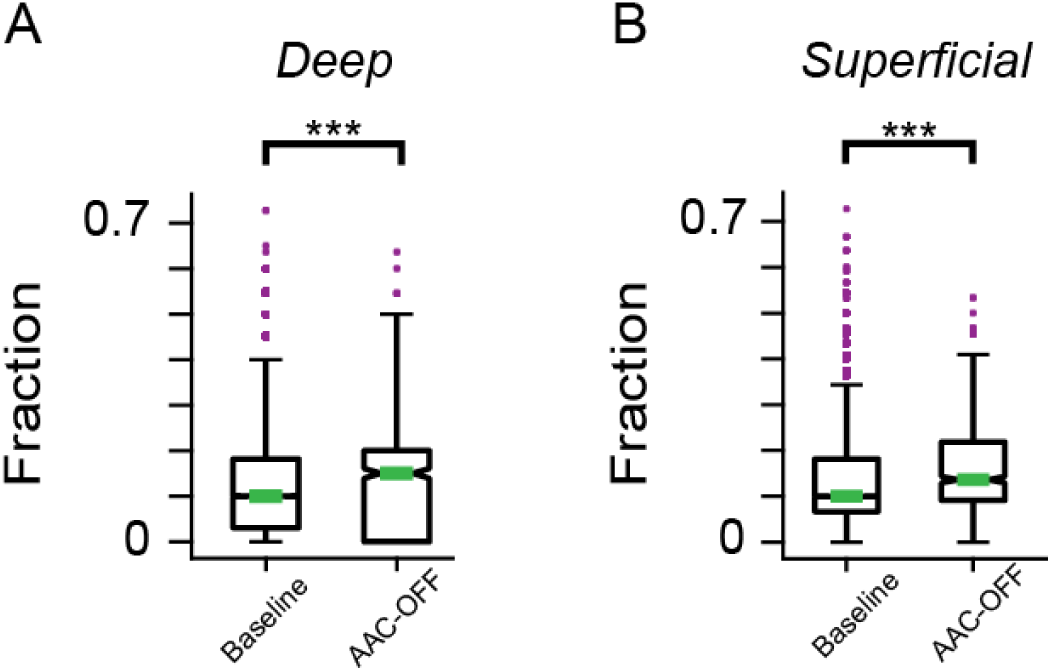
Both the fraction of deep and superficial PYRs participating is increased in AAC-OFF SPW-R. PYR cell participation separated by soma depth (n=1,495-12,364, total: 37,032 Control SPW-R. n=51-287, total: 868 AAC-OFF SPW-R.) A.) Fraction of Deep PYR participating in SPW-R (n=11-32, total: 125 Deep cells, p= 2.4645e-13, Wilcoxon rank-sum test). B.) Fraction of Superficial PYR participating in SPW-R (n=5-33, total: 69 Superficial cells. p= 6.5141e-06, Wilcoxon rank-sum test).

